# In vivo directed evolution of AAV in the primate retina

**DOI:** 10.1101/847459

**Authors:** Leah C. Byrne, Timothy P. Day, Meike Visel, Cecile Fortuny, Deniz Dalkara, William H. Merigan, David V. Schaffer, John G. Flannery

**Affiliations:** Helen Wills Neuroscience Institute, University of California Berkeley; INSERM U968, Institut de la Vision, 75012 Paris, France; UMRS968, Institut de la Vision, Sorbonne Universités, Pierre et Marie Curie University (UPMC) University Paris 06, Centre National de la Recherche Scientifique (CNRS) UMR7210, Institut de la Vision, 75012 Paris, France; Flaum Eye Institute, University of Rochester

**Author notes:** Correspondence should be sent to David Schaffer (278 Stanley Hall, Berkeley, CA 94720-3220) or John Flannery (132 Barker Hall, Berkeley, CA 94720-3190). David Schaffer and John Flannery participated equally. Departments of Ophthalmology, Neurobiology and Bioengineering, University of Pittsburgh (Leah C. Byrne).

## Abstract

Efficient AAV-mediated gene delivery remains a significant obstacle to effective retinal gene therapies. Here, we apply directed evolution – guided by deep sequencing and followed by direct *in vivo* secondary selection of high-performing vectors with a GFP-barcoded library – to create AAV viral capsids with new capabilities to deliver genes to the outer retina in primates. A replication incompetent library, produced via providing *rep* in trans, was created to mitigate risk of AAV propagation. Six rounds of *in vivo* selection with this library in primates – involving intravitreal library administration, recovery of genomes from outer retina, and extensive next generation sequencing of each round – resulted in vectors with redirected tropism to the outer retina and increased gene delivery efficiency to retinal cells. These new viral vectors expand the toolbox of vectors available for primate retina, and may enable less invasive delivery of therapeutic genes to patients, potentially offering retina-wide infection at a similar dosage to vectors currently in clinical use.

## Introduction

Inherited retinal degenerations (RDs) are caused by mutations in >200 genes(*1*), the majority of which are expressed in photoreceptors or retinal pigment epithelium (RPE) of the outer retina, leading to vision loss and decreased quality of life(*2, 3*). Gene therapy is a promising approach to treating RDs, as evident in the recent FDA approval of a gene therapy to treat inherited retinal degenerations with biallelic RPE65 mutations. While administration of a therapy to the vitreous fluid of the eye offers a simple and less invasive route compared to the subretinal surgery used for RPE65 and other treatments, as well as the potential to transduce the entire retina, efficient gene delivery to the outer retina in general remains a significant hurdle(*4*). Directed evolution has enabled the creation of new, efficient viral vectors for outer retinal gene delivery in mice(*5-7*). However, the anatomical features and structural barriers of large animal retinas are substantially different from mouse(*8*), as for example large animals have a specialized area for high acuity vision (the area centralis in dogs or fovea in primates), a vitreous of thicker consistency, and a thicker inner limiting membrane(*9, 10*). As a result, vectors that were selected for delivery in mouse retina are not as efficient in large animal models as in rodents (*7*).

Here, we used directed evolution(*11*), guided by insights into the population dynamics of viral selections that were derived from deep sequencing, to create AAVs capable of delivering genes to outer retina following intravitreal injection in primates, which have eye structures similar to humans. Subsequent, secondary screening of subsets of the resulting viral variants revealed the most efficient AAVs for photoreceptors and RPE of the primate retina. The outer retinal gene delivery offered by these variants may enable less invasive delivery of therapeutic genes to the retina, and potentially provide retina-wide transfection at a similar dosage.

## Results

### Directed evolution of AAV in primate retina

The nonhuman primate is a critical preclinical model for human therapeutic development, as it has a retinal anatomy similar to that of humans. In particular, primates are the only large animal model that possess a fovea, the specialized high acuity area of the retina. The fovea is essential for daily activities such as reading, is critical to quality of life, and is lost in numerous retinal degenerations. Here, we have conducted AAV directed evolution in a primate model. Our previous efforts in mouse were based on replication-competent AAV genome libraries; however, primates are known to often harbor a co-infection of herpes B (a helper virus for AAV). In order to mitigate the possibility of replication of the AAV libraries in the primate retina and subsequent spread, a *rep* in trans strategy was developed in which the library *rep* sequence is mutated (pRepSafeStop), while leaving intact regulatory elements, including the P40 promoter, which is necessary for *cap* protein expression. Stop codons in the *rep* sequence prevented the expression of the four Rep proteins, and the full *rep* sequence was supplied on pRepIntronHelper. This system resulted in a greater than 10-fold reduction in replication in the presence of high titers of adenovirus. The ITRs in AAV are highly recombinogenic, leading to the possibility of recombination and subsequent replication in the presence of a helper virus when using this strategy, so an intron was inserted in the *rep* supplied in trans to prevent packaging of recombined genomes (Figure 1). Replication incompetent libraries were constructed, packaged, and included in the primate screen including AAV2-7mer, Ancestral-7mer(*12*) and LoopSwap(*13*) libraries.

**Figure 1.**
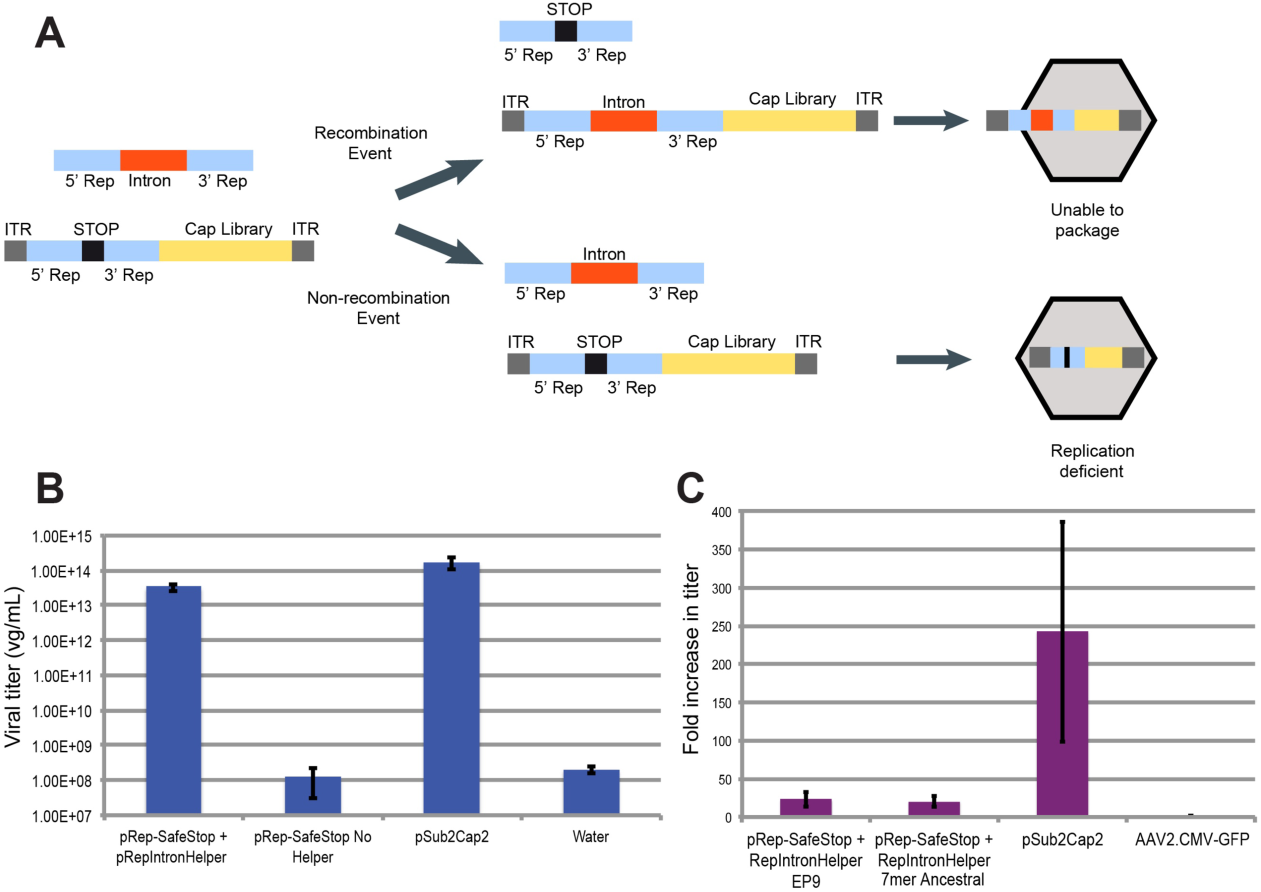
Replication incompetent AAV libraries. (**A)** Schematic depicting both recombination and non-recombination events with the pRepIntronHelper. If a recombination event were to occur, the intron sequence (*nebulin* intron 8, 774 bp) would push the transgene over the packaging capacity of AAV, leading to incomplete packaging. If recombination does not occur, the mutated *rep* sequence will be packaged, mitigating the possibility of replication. **(B)** Titering of the *rep* in trans system with and without pRepIntronHelper, in comparison to the transgene with native *rep*, pSub2Cap2. The *rep* in trans system leads to similar titers as normal pSub2Cap2 packaging. **(C)** An adenovirus rescue study determined that the *rep* in trans system leads to greater than 10-fold reduction in replication. The abilities of an AAV9 error prone library and the 7mer-Ancestral library to replicate with the *rep* in trans system are shown, compared to an AAV2 with the wild-type genome and a replication incompetent AAV2 with a CMV-GFP transgene.

Libraries were injected, harvested, and repackaged for up to 5 sequential rounds of selection, with one round of error prone PCR performed after round 3 (Figure 2, Supplemental Table 1). AAV *cap* genes were PCR amplified from the outer nuclear layer (ONL), which was isolated from transverse cryosections of retina, and in parallel from separated RPE (Supplemental Figure 1).

**Figure 2.**
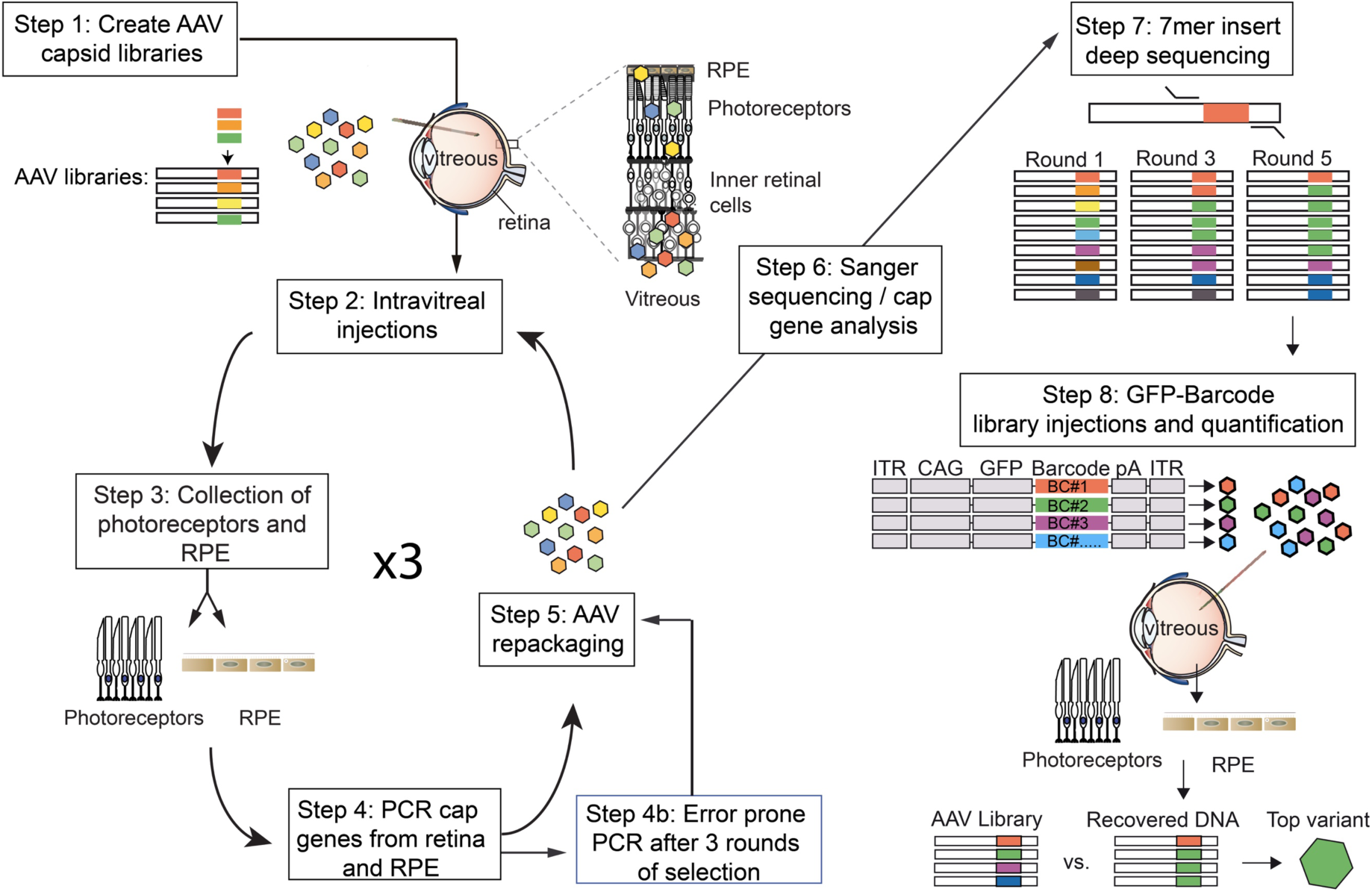
Workflow of directed evolution of AAV in the primate retina. Highly diverse (∼1E+7) libraries of AAV variants were packaged such that each virus contained a genome encoding its own capsid. Libraries were pooled and injected intravitreally in primates. After AAV infection had occurred, retinal tissue and RPE cells were collected, and *cap* gene variants were PCR amplified, recloned, and repackaged for the subsequent round of injection. Five rounds of selection were performed, and error prone PCR was performed after the third round to introduce additional diversity into the library. Following the selections, each pool was subjected to deep sequencing to analyze the dynamics of each individual variant and overall convergence of the library. Based on their increase in representation relative to the original library, individual variant capsids were chosen and used to package a scCAG-eGFP genome also containing a unique DNA barcode sequence. These barcoded vectors were then pooled in equal amounts and injected intravitreally. Retinal cells (photoreceptors or RPE cells) were harvested, GFP barcodes were PCR amplified from the collected tissue, and deep sequencing was used to quantify the relative abundance of barcodes. The top-performing variants were evident as those with the greatest fold increase of barcodes recovered from collected tissue relative to the injected library.

Deep sequencing (source data 1) revealed that libraries contained ∼1E+6 - ∼1E+7 individual variants, which converged to ∼1E+4 - ∼1E+5 variants over 6 rounds of selection, a diversity not possible to observe through Sanger sequencing (Figure 3A). In each of the libraries analyzed, a small portion of library members were originally over-represented in the initial plasmid library (Figure 3B). However, relative to this input, analysis of results from deep sequencing over the rounds of selection revealed a subset of variants that increased significantly in their representation during rounds of selection for each of the input libraries (Figure 3C).

**Figure 3.**
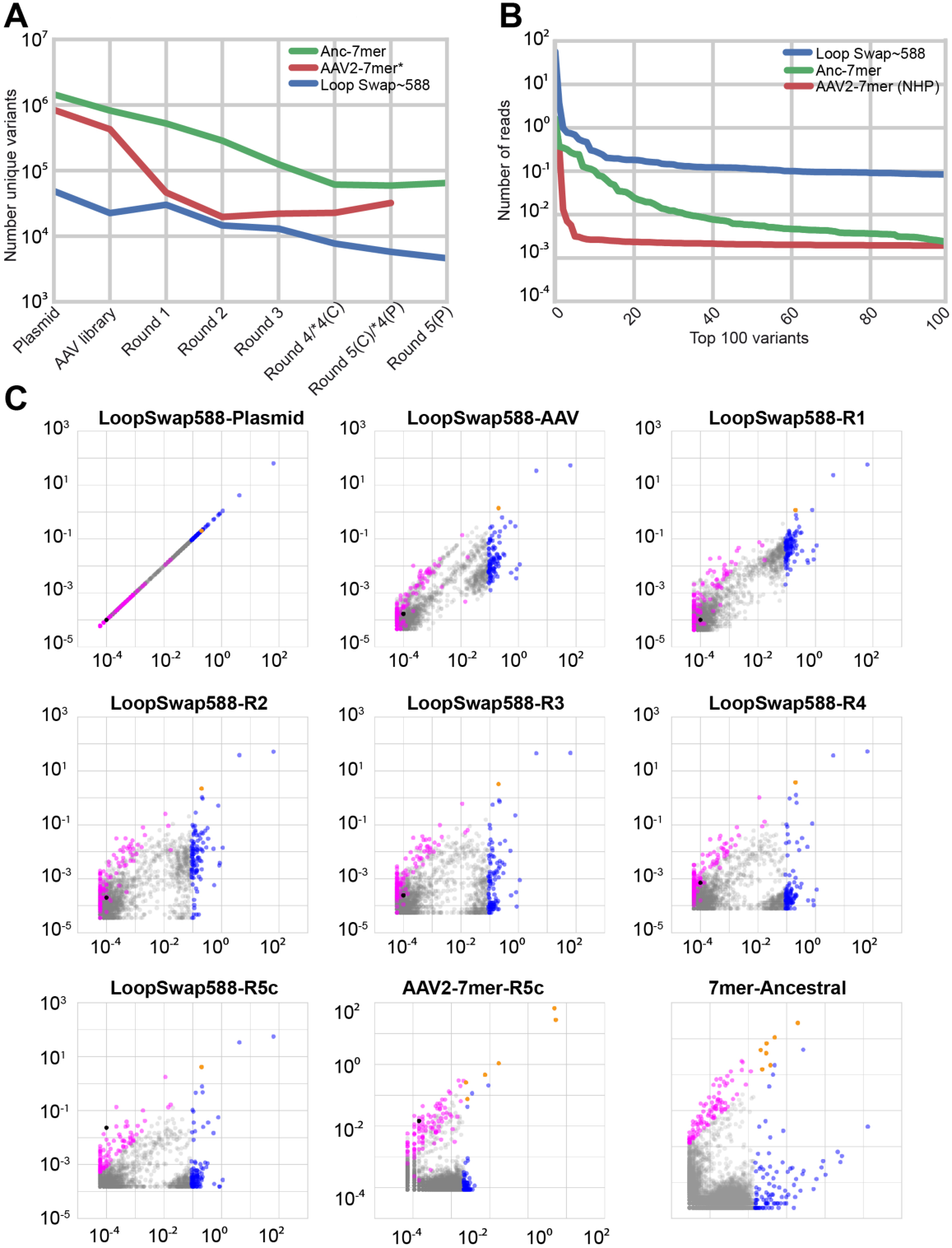
Directed evolution of AAV in primate retina. **(A)** Deep sequencing of libraries revealed convergence of variants over rounds of selection. **(B)** In each of the libraries evaluated, a small proportion of variants were overrepresented in the plasmid library. **(C)** Scatterplots illustrate the behavior of individual variants over all rounds of selection for the ∼588 LoopSwap library for all rounds of selection, and at the final round of selection for AAV2-7mer and 7mer-Ancestral libraries. Additional scatter plots are shown in Supplementary Figure 2. Black dots in the LoopSwap plots indicate variant NHP#26, validated in Figure 6. The black dot in the AAV2-7mer plot indicates variant NHP#9, validated in Figure 5. A pseudo-count of 1 was added to each variant prior to plotting. X-axis is the percent of the library made up by each variant in the original library. Y-axis is the percent of total library at the indicated round of selection. As variants increase in representation they rise on the Y-axis. Variants overrepresented in the original library are colored blue. Variants that had the greatest fold increase in representation in the final round of selection are shown in magenta. Variants that were overrepresented in the original library and increased significantly in representation over rounds of selection are colored orange. From the last round of selection, sequencing was performed on samples from central (R5C, Supplemental Figure 2) and peripheral (R5P) samples separately.

### Secondary barcoded-GFP library screening in primate retina

Twelve variants from the three successfully amplified libraries were chosen for a secondary round of selection with GFP-barcoded libraries, along with parental serotypes as controls. Specifically, each capsid was used individually to package a recombinant genome containing GFP plus a 3’ barcode unique to that capsid, and the resulting vectors were pooled at equal ratios. This new library was injected in both eyes of a primate, and 3 weeks after injection, biopsies were collected from locations across the retina (Figure 4A). GFP expression resulting from injection of the GFP-barcode libraries was primarily found in photoreceptors, as well as some inner retinal cells, a tropism that is shifted from AAV2 or 7m8, which yielded stronger inner retinal expression(*7*) (Figure 4B). The outer nuclear layer and RPE were anatomically isolated, DNA was purified from these samples, and deep sequencing was performed to quantify the relative extents to which each capsid was capable of delivering its genome deep into the retina from the vitreous.

**Figure 4.**
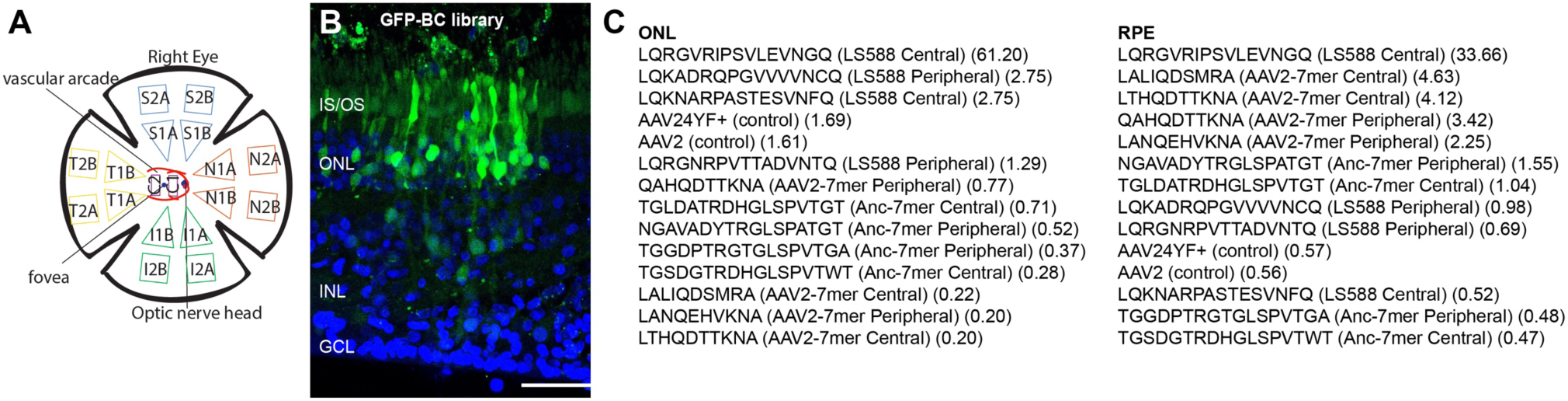
Directed evolution of AAV in primate retina. **(A)** A map of the primate retina shows the distribution of samples that were collected for rounds of selection and the GFP-barcode library. **(B)** GFP expression (shown here from retina along the superior vascular arcade) resulting from the barcoded library revealed that expression was shifted to an outer retinal tropism in selected variants. Scale bar is 50 µm. **(C)** GFP-barcode library injection results, for primate outer retina. The lists of variants are ordered from best (top) to worst (bottom) performing vectors, along with a description of the source library and the sample the variant was identified from (central or peripheral) and a value indicating the extent to which the variant competed with other vectors, expressed as: % of total in recovered library/% of total in AAV library.

### Validation of the top-performing primate variants

Quantification of vector performance in the ONL revealed that AAV2-7mer and Loop swap based variants outperformed other viruses (Figure 4C). The top-ranking vector, Loop Swap variant AAV2 583∼LQRGVRIPSVLEVNGQ, outperformed other variants in the GFP-barcode screen, though it yielded lower viral titers (∼5E+11 vg/mL). AAV2-LALIQDSMRA (designated NHP#9), the second ranking variant from the GFP-barcode screen in RPE, packaged at high titers (∼5E+13 vg/mL) and was therefore selected for a first round of validation studies focusing on ganglion cells of the inner retina and cones of the outer retina. Cone photoreceptors are involved in adult macular degeneration (AMD), which is the most common cause of blindness in developed countries and is predicted to affect 288 million people worldwide by the year 2040(*14*), and are therefore a primary target for retinal gene therapy.

NHP#9 and the previously described murine variant 7m8(*7*) were packaged with a gamma-synuclein gene (SNCG) promoter to drive tdTomato expression in RGCs(*15*) and the pR1.7(*16*) promoter to yield GFP expression in cones. Vectors encoding both these constructs were mixed in equal ratios (∼1.5E+12 vg/construct/eye) and injected intravitreally in a cynomolgous monkey. Expression of tdTomato in RGC’s was lower in NHP#9-injected eyes compared to 7m8, which infected ganglion cells across the expanse of the retina efficiently (Figure 5); however, expression in foveal cones was increased relative to 7m8, indicating a shift in tropism away from the inner retina towards photoreceptors in the outer retina. Changes in the efficiency of expression following injection of 7m8 and NHP#9 were evaluated by two methods: the numbers of RGC’s and cones infected were quantified by imaging, and qRTPCR was used to quantify levels of expression in these cells. Quantification of cell numbers, performed using Imaris software on confocal images from the macula, revealed that 7m8 infected 10,310 RGC’s, while NHP#9 infected 4,296. In contrast, NHP#9 infected 2202 cones, compared to 1019 cones infected by 7m8 (Figure 5G,H). qRT-PCR, performed using the ΔΔCT method, revealed an 11.71 (10.37 - 13.22) fold increase of GFP expression in foveal cones relative to 7m8.

**Figure 5.**
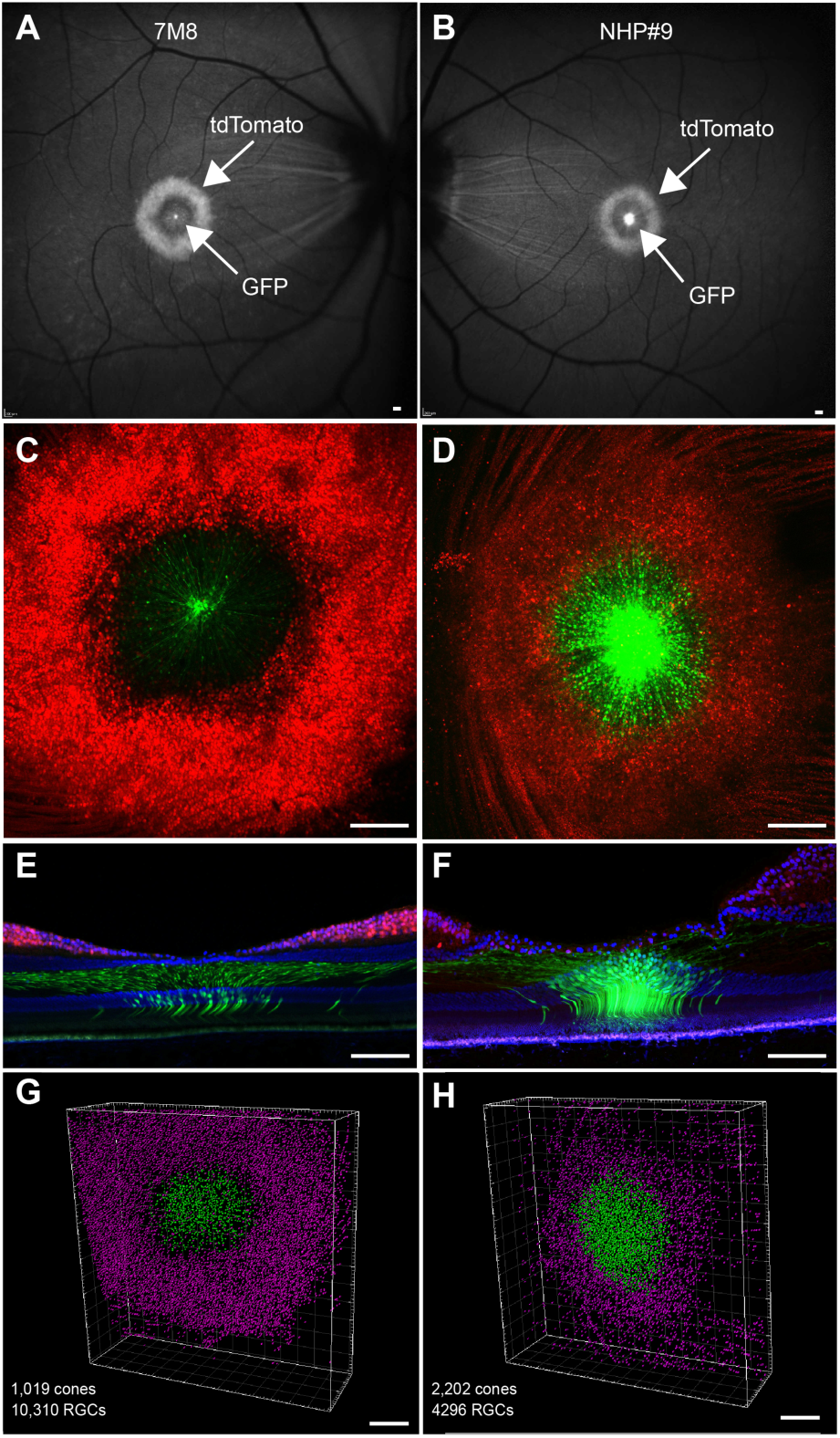
Validation of NHP#9 in primate retina. (**A-H**) Co-injection of ∼1.5E+12 particles of SNCG-tdTomato and ∼1.5E+12 pR1.7-eGFP packaged in 7m8 and variant NHP#9 in primate retina. Intravitreal injection of 7m8 **(A**,**C**,**E)** resulted in robust tdTomato expression in ganglion cells and expression of GFP in foveal cones. In contrast, injection of equal number of particles of NHP#9 in the contralateral eye resulted in reduced ganglion cell expression, and increased GFP expression in cones relative to 7m8 **(B**,**D**,**F). (G**,**H)** Quantification of ganglion cells and cones transduced with 7m8 and NHP#9 in primate retina. Counting of labeled cells, performed using Imaris software, revealed a substantial decrease in numbers of transduced ganglion cells and an increase in the number of cones targeted with NHP#9, compared to 7m8. Scale bars in A,B: 200 µm, C-F: 100 µm, G,H: 200 µm.

The top-ranking variant from the GFP barcode screen, Loopswap variant ∼583-LQRGVRIPSVLEVNGQ (designated NHP#26), was also tested for validation despite the limitation that reduced production of this vector enabled only a low dose. ∼5E+10 particles of NHP#26-scCAG-eGFP were injected intravitreally into one eye of a cynomolgous monkey. Although the number of particles injected was 30-fold lower than for the other tested vectors, efficient expression of GFP was observed in the fovea and in regions across the retina (Figure 6). In contrast to the foveal-spot-and-ring pattern of expression that was observed with 7m8, NHP#9, and other naturally occurring serotypes, imaging within the foveal region of NHP#26 resulted in a disc of GFP expression centered on the foveola (Figure 6A). Confocal imaging of the flatmounted retina confirmed this disc pattern of expression around the fovea (Figure 6B), with very few GFP positive ganglion cell axons. Punctate regions of GFP expression were often strongest around retinal blood vessels (Figure 6C), and were located across the expanse of the retina. Imaging of cryostat sections taken from the retina confirmed that there was little GFP expression in ganglion cells, as indicated by the lack of GFP+ ganglion cell axons, while high levels of GFP expression were found in Müller cells, additional cells in the inner nuclear layer, some foveal cones, and rods across the retina (Figure 6D-K).

**Figure 6.**
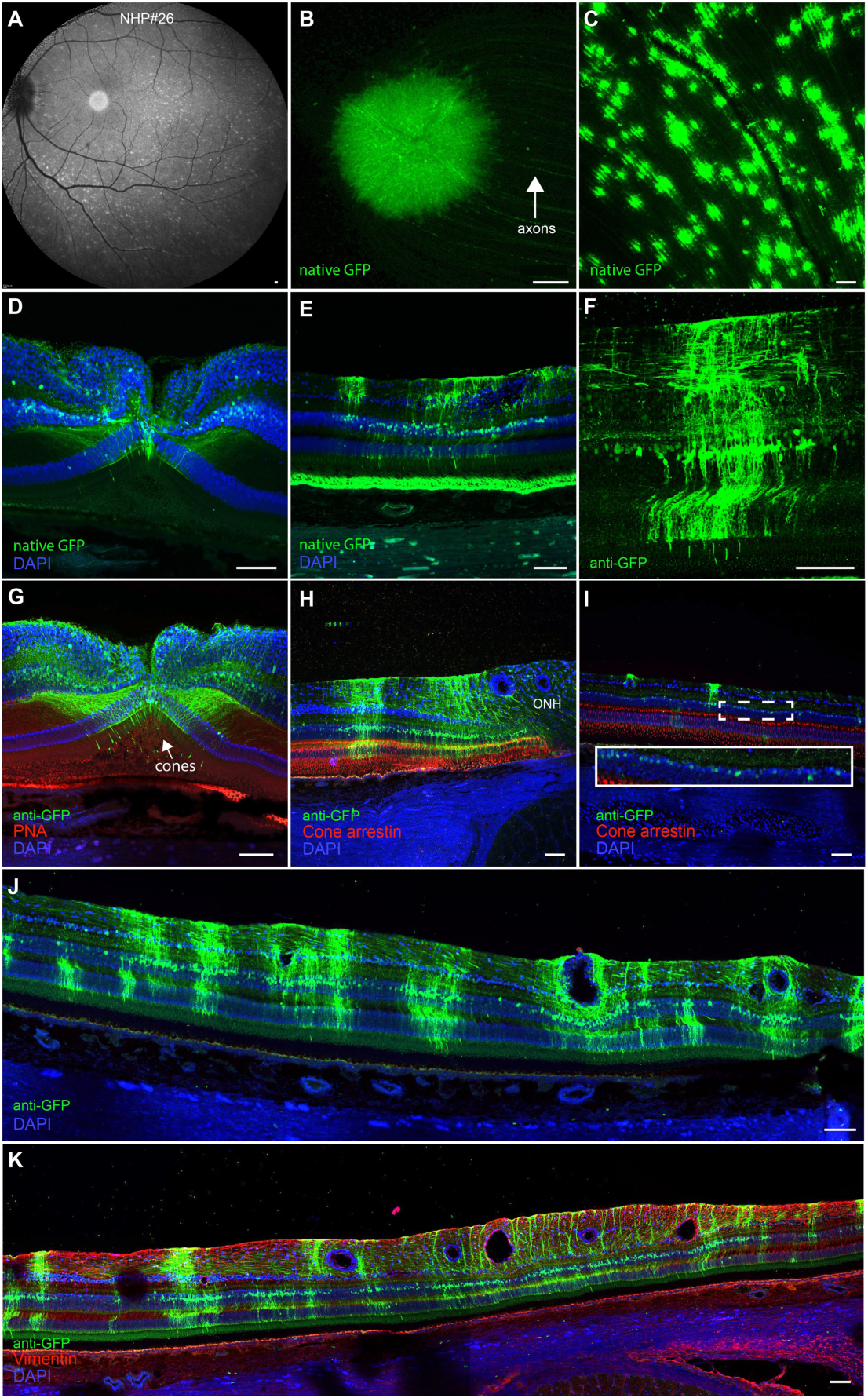
Validation of NHP#26 in primate retina. **(A)** Fundus imaging in a primate eye following injection of 5E+10 particles of NHP#26-scCAG-GFP revealed a disc of GFP expression centered on the fovea, and a punctate pattern of GFP expression across the retina. **(B)** Confocal imaging of native GFP expression in the flatmounted fovea. **(C)** Confocal imaging of native GFP expression in the area outside of the vascular arcade. **(D)** Confocal imaging of native GFP expression in a cryostat section through the fovea. **(E)** Native GFP expression in the inferior retina, outside the vascular arcade, shows little GFP expression in ganglion cells, but high levels of expression in Müller cells and some photoreceptors in the outer retina. Autofluorescence was also observed in RPE. **(F)** Anti-GFP labeling in a cryostat section revealed GFP expression in photoreceptors, evident by their outer segments, Müller cells, evident by their retina-spanning processes, as well as cells in the inner nuclear layer with horizontal processes that are likely interneurons. **(G)** Anti-GFP labeling in a foveal section reveals additional infected cones, Müller glia, and interneurons. **(H)** Co-labeling with anti-cone arrestin and anti-GFP antibodies reveals GFP expression in rod photoreceptors, as well as cells in the inner nuclear layer, in a section taken next to the optic nerve head. **(I)** Co-labeling with anti-cone arrestin and anti-GFP antibodies in an area of low expression reveals GFP expression in inner nuclear layer cells. **(J**,**K)** Montages of confocal images from cryostat sections collected outside the vascular arcade show efficient expression of GFP in the inner nuclear layer and outer retina. Scale bars in A,B: 200 µm, C-K: 100 µm.

## Discussion

Clinical trials using subretinal surgeries have shown the promise of AAV-mediated gene therapy for retinal disease(*17, 18*); however, efficient delivery of therapeutic genes across the outer retina – such as from an intravitreal injection – may substantially enhance the safety and retinal area of treatment of gene therapies in such patients. Together, the results described here show that directed evolution, guided by deep sequencing, enabled identification of novel AAV viruses that were not observable by Sanger sequencing and had an enhanced ability to infect cells in the outer retina of primates, an important large preclinical animal model for the development of retinal therapies.

Deep sequencing to gain insights into selection dynamics revealed that the variants with greatest fold increase during selection, rather than the most frequent final variant, is the optimal metric for identifying top-performing variants. Overrepresentation of variants in the original library significantly influenced the number of clones in the final round of selection, with variants that were highly represented in the original library more likely to constitute a majority of clones in the final round, and masking more promising variants that were less abundant at the start but that had a greater fold increase over rounds of selection.

Also, the use of GFP-barcoded libraries enabled the selection of, from a pool of top-performing candidates identified from deep sequencing, AAV variants with best transduction efficiency for the targeted cells. Barcoded library screening represents a sensitive method for evaluating many variants in parallel, in the same animal, allowing for direct, head-to-head comparison and thus reducing animal numbers(*19*). Development of vectors in preclinical models must continue to take into account the anatomical differences that exist between species, and the possibility that optimization of a therapy in small animals may not translate into large animals with anatomies more similar to humans(*20*).

The primate-derived variant NHP#26 transduced the outer retina at greatly reduced titer. Increasingly, it is understood that immune response to high doses of AAV vectors represents a significant factor influencing the success of retinal and other gene therapies(*21, 22*). This vector may enable safer gene therapies for retinal degeneration in patients as a result of the decreased vector load required for transgene expression and may reduce risk of vector-related toxicity or immunogenicity.

Additional studies into the composition of physical barriers, including the vitreous and inner limiting membrane, may elucidate the physical basis for the patterns of expression seen following intravitreal injection. Furthermore, directed evolution screens isolating variants from specific retinal locations, such as the macula, may result in variants with increased capabilities to target these areas, as results from the present screens represent an average across the retina. Deeper characterization of the vectors created in this study will likely lead to additional mechanistic insights into cell targeting and tropism.

## Methods

### Construction of the pRepSafeStop directed evolution backbone

The pRepSafeStop plasmid containing a *Not* I site for *cap* cloning was created by Quikchange site-directed mutagenesis(SDM) on pSub2Cap2^14^ to introduce stop codons in *rep* at codons 5 and 235 using pRepSafeStop SDM primers. Unique pRepSafeStop backbones containing *Asc* I and *Spe* I sites were created via Gibson Assembly in order to maintain separation of libraries through rounds of selection. Libraries were PCR amplified and digested with *Hind* III and *Not* I/*Asc* I/*Spe* I, then ligated into the pRepSafeStop construct.

### Rep intron helper cloning

Intron 8 from the *nebulin* gene was amplified from genomic DNA isolated from HEK293T cells using the Neb Genomic primers and cloned into a TOPO vector. To create the pRepIntronHelper, the rh10 AAV helper plasmid pAAV2/rh10 was digested with *Pme* I and *Bsm* I, Klenow blunting was performed, and the resulting DNA fragment was re-circularized. The first AGG sequence after the Rep52/Rep40 start codon was chosen as the site for intron insertion based on a computational analysis of splice signal motifs. Infusion assembly (IFA) was used to insert the *nebulin* intron using the IFA primers, thereby generating the final plasmid.

### Adenovirus rescue to determine loss of AAV replication with the pRepSafeStop backbone

HEK293T cells were infected with AAV (AAV2, an EP9 variant and a 7mer Ancestral variant) at an MOI of 10^5^ and then incubated at 37° C and 5% CO_2_ for 48 hours. Next, 10 µL of an adenovirus 5 (Ad5) stock, a volume that resulted in cytopathic effect within 24 hours, was added, and the plates were incubated at 37° C and 5% CO_2_ for an additional 48 hours. Cells that were then harvested, pelleted, resuspended in 100 µL of lysis buffer (0.15M NaCl, 50 mM Tris HCl, 0.05% Tween, pH 8.5). Freeze/thaws were used to lyse the cells, and 5 µL of the crude lysate was used for titering AAV to quantify replication.

### AAV selection in primate outer retina

Three weeks after intravitreal injection, the primate was euthanized, and both eyes, as whole globes, were briefly submerged in 4% paraformaldehyde. Superior, inferior, temporal, and nasal regions of the retina were cut into pieces, and the RPE was separated from each section. Retinal sections were then immersed in 30% sucrose, embedded in OCT media, and flash frozen. Retinal pieces were sectioned transversely at 20 µm. During sectioning, DAPI staining and light microscopy were used to identify each nuclear layer in the retina, and the inner nuclear and ganglion cell layers were removed. DNA was extracted from samples using a Qiagen DNeasy blood and tissue kit, according to manufacturer’s instructions.

### AAV packaging

AAV libraries were constructed prior, and have been previously described^7,8,15,16^. After each round of injection, capsid sequences were recovered by PCR from harvested cells using primers HindIII_F1 and NotI_R1, AscI_R1, or SpeI_R1, with reverse primers being specific to unique AAV backbones, in order to maintain separation of groups of libraries. PCR amplicons were then digested, and recloned into the AAV pRepSafeStop backbone. AAV packaging has been described previously^15,17^. AAV vectors with pRepSafeStop backbone were produced by triple transient transfection of HEK293T cells with the addition of the pRepIntronHelper plasmid in 5 times greater concentration than the library plasmid, purified via iodixanol density centrifugation, and buffer exchanged into PBS by Amicon filtration. DNase-resistant viral genomic titers were measured by quantitative real time PCR using a BioRad iCycler.

### Deep sequencing of AAV libraries from rounds of selection

A ∼75-85 base pair region containing the 7mer insertion or Loop Swap mutation sites (semi-random mutations at surface exposed regions, for a description of Loop Swap library construction see (*13*)) was PCR amplified from harvested DNA. Primers included Illumina adapter sequences containing unique barcodes to allow for multiplexing of amplicons from multiple rounds of selection (Supplemental Table 2). PCR amplicons were purified and sequenced with a 100-cycle single-read run on an Illumina HiSeq 2500. Custom Python code was written for analysis. First, the DNA sequences encoding amino acid insertions, found between constant linker DNA sequences were identified. Then, DNA sequences were translated into amino acid sequences. The number of reads for each amino acid insertion sequence was then counted, across the AAV library and across rounds of selection. Read counts were normalized by the total number of reads in the run. Pandas (https://pandas.pydata.org) was used to analyze dynamics of directed evolution and create plots.

### Deep sequencing analysis

Deep sequencing was performed at >10X depth of the number unique variants in the round. Reads with low quality scores were eliminated from further analysis using Illumina workflow. Variants were analyzed on the amino acid level (i.e. variants with varying DNA sequences encoding the same amino acid sequence were pooled together for analysis). Best performing variants were chosen as variants with the greatest fold increase in the final round of selection relative to the initial plasmid library (# reads in final round, normalized to total number of reads in the round / # of reads in library, normalized to total number of reads in the round). A pseudo-count of 1 was added before normalization to each individual variant to allow analysis of variants not appearing in sequencing of the plasmid library(*23*).

### GFP barcode library construction

Unique 25 bp DNA barcodes were cloned behind a self-complementary AAV ITR construct containing a CAG promoter driving eGFP (CAG-GFP-Barcode-pA). Individual variants were packaged separately with constructs containing different barcodes. Variants were then titer matched and mixed in equal ratios before injection into primates.

### Deep sequencing of GFP-barcode libraries

Barcodes were PCR amplified directly from DNA which was harvested from primate retinal tissue. Samples were collected from areas across the retina, and from ONL or RPE. Primers amplified a ∼50 bp region surrounding the GFP barcode and contained Illumina adapter sequences and secondary barcodes to allow for multiplexing of multiple samples (Supplemental Table 2). PCR amplicons were purified and sequenced with a 100-cycle single-read run on a MiSeq. Read counts were normalized by total number of reads in the run. Analysis of barcode abundance was performed using custom code written in Python, followed by creation of plots in Pandas. Barcodes were identified as variable DNA sequences found between constant sequences in the expression cassette, which surrounded the barcode. Best performing variants were selected based on the fold increase in the percent of total library, relative to the injected library (% of total in recovered sample / % of total in injected library). Analysis was performed on n=1 primate.

### Gene expression analysis

GFP expression was calculated relative to GAPDH, using the ΔΔCT method on RNA collected from retinal sections and extracted using AllPrep DNA/RNA FFPE Kit (Qiagen). Tests were performed in triplicate with technical replicates from the same eye.

### Primers

Primer sequences are listed in Supplemental Table 2.

### Animal studies

#### Mice

C57BL/6 mice from Jackson Laboratories were used for mouse experiments. Surgery was performed under anesthesia, and all efforts were made to minimize suffering.

#### Primates

Cynomolgous monkeys between 4-10 years old were used for all studies, and intravitreal injections were made with methods described previously(*24*). In order to prevent any immune response to GFP or viral vectors, which have been previously reported *(7)*, the monkey used for fluorophore expression received daily subcutaneous injections of cyclosporine at a dose of 6 mg/kg for immune suppression, beginning one week before AAV injection, and adjusted based on blood trough levels to within a 150-200 ng/ml target range. All primates were screened for neutralizing antibodies prior to inclusion in the study and had titers <1:25. Confocal scanning laser ophthalmoscopic images (Spectralis HRA, Heidelberg Engineering) were obtained 3 weeks after injection, with autofluorescence settings, which lead to effective tdTomato and GFP visualization. For histology, the monkey was euthanized, both retinas were lightly fixed in 4% paraformaldehyde, and tissue was examined by confocal microscopy. At the conclusion of the experiment, euthanasia was achieved by administering an IV overdose of sodium pentobarbital (75 mg kg-1), as recommended by the Panel on Euthanasia of the American Veterinary Medical Association. Pieces of primate retina were then prepared in 30% sucrose, embedded in OCT media, flash frozen, and sectioned at 20 µm for confocal microscopy imaging of native fluorophore expression. Antibodies for labeling were: anti-GFP (A11122, Thermo, 1:250) anti-vimentin (Dako, 1:1000), peanut agglutinin (PNA, Molecular Probes, 1:200), and anti-cone arrestin (7G6, gift from Peter MacLeish, 1:50). A summary of minor adverse events related to the procedures is summarized in Supplemental Table 1.

## Study Approval

### Mice

All procedures were performed in accordance with the ARVO statement for the Use of Animals in Ophthalmic and Vision Research, and were approved by the University of California Animal Care and Use Committee AUP# R200-0913BC.

### Primates

The procedures were conducted according to the ARVO Statement for the Use of Animals and the guidelines and under approval from of the Office of Laboratory Animal Care at the University of Rochester.

## Author contributions

LCB: Conceived, planned and executed experiments. Analyzed data. Wrote the manuscript. TPD: Conceived, planned and executed primate experiments. Analyzed data. Wrote the manuscript. MV: Planned experiments, performed AAV packaging, analyzed data. Wrote the manuscript. CF: Conceived, planned and executed primate experiments. DD: Performed directed evolution screening in macaque retina, provided AAV constructs with PR1.7 / SNCG promoters and wrote the manuscript. WHM: Supervised intravitreal injection and fluorescent fundus imaging of viral vectors in macaques. Wrote the manuscript. DVS: Conceived, planned and supervised project. Wrote the manuscript. JGF: Conceived, planned and supervised project. Wrote the manuscript.

## Acknowledgments

Deep sequencing was performed at the UC Berkeley Vincent J. Coates sequencing facility. Confocal imaging was performed at the Berkeley Biological Imaging Facility. Injections for AAV selection in primates were performed at Valley Biosystems. We thank Yvonne Lin and Jaskiran Mann for technical assistance.

