## Supplemental materials for "In vivo directed evolution of AAV in the primate retina"

### Supplementary Materials

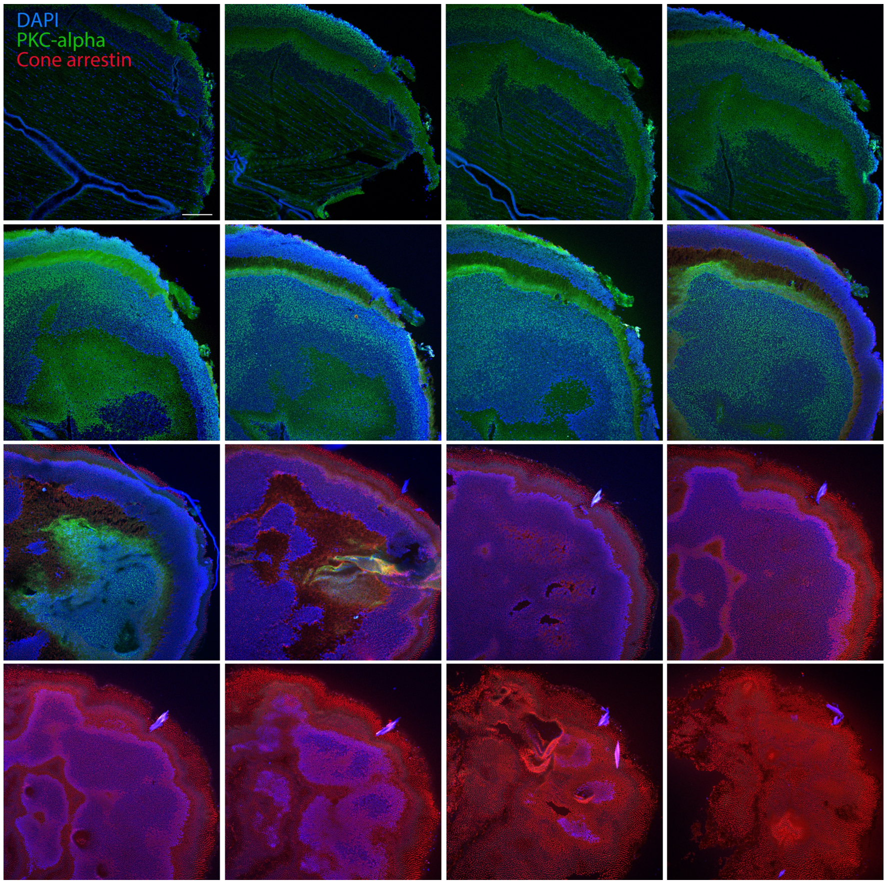


**Supplementary Figure 1. Transverse cryosectioning of mouse retinal punch illustrates the method used to isolate the outer retina.** A punch of retina was flatmounted, embedded in OCT cutting medium, and flash frozen, then mounted in a cryostat and sectioned at 20 µm sections through retinal layers. Layers were then stained for PKC-alpha (a marker of bipolar cells in the inner nuclear layer) and cone arrestin (a marker of photoreceptors in the outer retina). Labeling shows successful isolation of outer retinal tissue from inner retinal cells, which was then used for amplification of libraries. RPE was peeled away prior to sectioning.

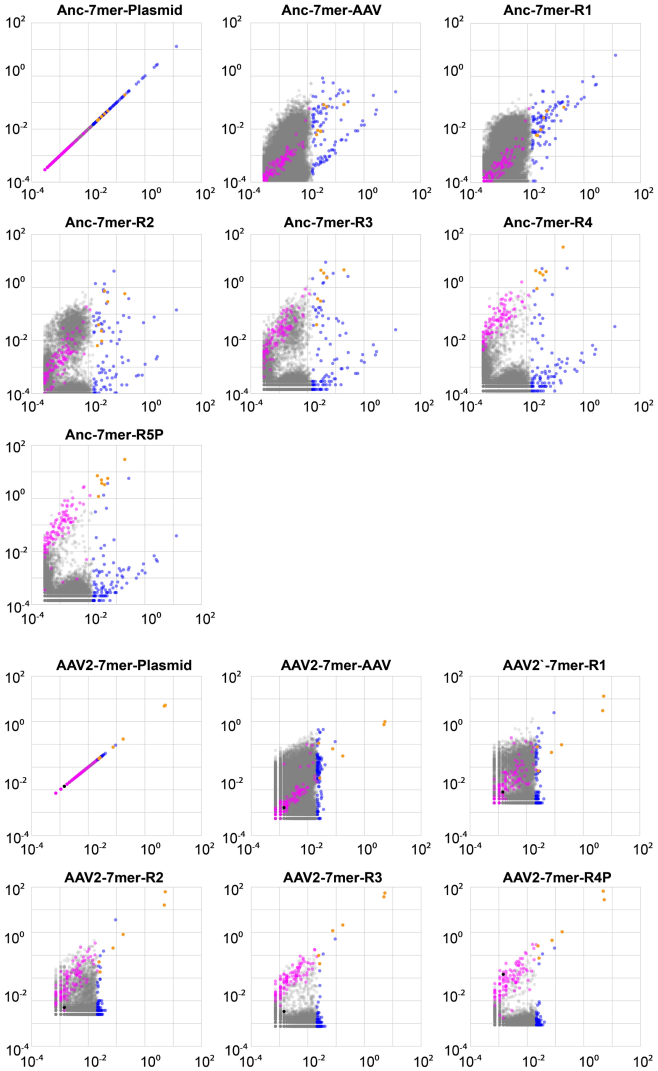


**Supplementary Figure 2. Scatter plots track variants across rounds of selection.** Scatterplots illustrate the behavior of individual variants over all rounds of selection for the ~Ancestral-7mer library and the 588-Loopswap library. Variants overrepresented in the original library are colored blue. Variants that had the greatest fold increase in representation in the final round of selection are shown in magenta. Variants that were overrepresented in the original library and increased significantly in representation over rounds of selection are colored orange. Black dots in AAV2-7mer scatter plots indicate the variant NHP#9.

| Round | NHP ID | Age/Weight/Sex | Libraries injected | Amount Virus Injected | Notes |
|---|---|---|---|---|---|
| 1 | V002278 | Approx. 7 years (age unknown at import)<br>5.98 kg/ Male | Loop Swap<br>Ancestral-7mer | 2.0E+11 vg per library; 100 µl volume. | |
| 2a | V002262 | Approx. 7 years (age unknown at import)<br>6.35 kg/ Male | Recovered variants from round 1<br>AAV2-7mer | 2.5E+11 vg ONL<br>2.5E+11 vg RPE<br>5.0E+10 vg AAV2-7mer; 100 µl volume. | No variants were PCR amplified following injection from this round, no obvious immune response noted. |
| 2b | V002148 | Approx. 7 years (age unknown at import)<br>6.48 kg/ Male | Recovered variants from round 1<br>AAV2-7mer | 1.3E+11 vg ONL<br>1.3E+11 vg RPE<br>5.0E+10 vg AAV2-7mer; 100 µl volume. | Repeat of previous round |
| 3 | V002265 | 8 years 5 months<br>4.92 kg/ Male | Recovered variants from round 3 | 4.3E+12 vg ONL<br>3.7E+12 vg RPE; 100 µl volume. | Error prone PCR conducted<br>No adverse events |
| 4 | V002540 | 6 years 9 months<br>4.59 kg/ Male | Recovered variants from round 4 | ~1E+12 vg per library; 100 µl volume. | No adverse events |
| 5 | V002861 | 6 years 6 months<br>6.60 kg/ Male | Recovered variants from round 5 | 2.4E+12 vg ONL<br>6.3E+12 vg RPE; 100 µl volume. | No adverse events |
| GFP-barcode | V002361 | 9 years 5 months<br>6.00 kg/ Male | Barcoded individual variants | ~1.0E+10 vg each variant; 100 µl volume. | Both eyes injected with GFP-BC library. Hyphema in left eye resolved in 12 days |
| Variant validation | 106 | 9 years 5 months<br>14.5 kg/ Male | 7m8 and NHP9 | ~1.5E+12 vg 7m8-pR1.7-GFP + 1.5E+12 vg 7m8-SNCG-tdTomato; 100 µl volume.<br>OR<br>~1.5E+12 vg NHP#9-pR1.7-GFP + 1.5E+12 vg NHP#9-SNCG-tdTomato; 100 µl volume. | No adverse events |
| Variant validation | 735 | 17 years<br>Male | NHP26 in one eye | ~5E+10 vg NHP#26-scCAG-GFP; 100 µl volume. | No adverse events |

**Supplemental Table 1**. Summary of the rounds of selection performed in primates. The table indicates the age and weight of the primates injected, the virus and titer injected at each round, and notes on the rounds of selection completed. ONL refers to virus libraries recovered from ONL samples. RPE refers to virus libraries recovered from RPE samples, which were processed in parallel. Round 2b was a repeat of the 2$^{nd}$ round of selection, which did not result in PCR amplification of variants.

| Primer | Sequence |
|---|---|
| SDM1 | GACCTTAATCACAATCTTTTAAAACCCCGGCATGGCGGCT |
| SDM2 | GGCTCGTGGACAAGTAAAGGGATTACCTCGGA |
| Neb Genomic_F | GTAAGGGTCTGCTCCATTGCCACTT |
| Neb Genomic_R | CTAAATCAAAAAAGAGTGAAAAGTTAGGAGG |
| IFA_F | TGGCTCGTGGACAAGGTAAGGGTCTGCTCCATTGC |
| IFA_R | CTCCGAGGTAATCCCCTAAATCAAAAAAGAGTGAAAAGTT |
| HindIII_F1 | GACGTCAGACGCGGAAGCTTC |
| NotI_R1 | GGTTTATTGATTAACAAGCGGCCG |
| AscI_R1 | TGGCGGACTTTATAGGCGCG |
| SpeI_R1 | GCCCAGTTCGAATAGCGAGT |
| LS588_Forward adapter | AATGATACGGCGACCACCGAGATCTACACTCTTTCCCTACACGACGCTCTTCCGATCT**NNNNNGTTCTGTATCTACCAACCTCCA** |
| LS588_rev_index1 | CAAGCAGAAGACGGCATACGAGAT**CGTGAT**GTGACTGGAGTTCAGACGTGTGCTCTTCCGATCTNNNNNGACCATGCCTGGAAGAACGCC |
| LS453_Forward adapter | AATGATACGGCGACCACCGAGATCTACACTCTTTCCCTACACGACGCTCTTCCGATCT**NNNNNATCGACCAGTACCTGTATTACT** |
| LS453_rev_index1 | CAAGCAGAAGACGGCATACGAGATCTGATCGTGACTGGAGTTCAGACGTGTGCTCTTCCGATCTNNNNNGGTCCCGAATGTCACTCGCTC |
| Anc_Forward adapter | AATGATACGGCGACCACCGAGATCTACACTCTTTCCCTACACGACGCTCTTCCGATCT**NNNNNCCTGCAGTCGTCTAACACCGC** |
| Anc_rev_index1 | CAAGCAGAAGACGGCATACGAGAT**CTCTAC**GTGACTGGAGTTCAGACGTGTGCTCTTCCGATCTNNNNNTAAGGCTCCCTGGCTGTTGAC |
| 5_Forward adapter | AATGATACGGCGACCACCGAGATCTACACTCTTTCCCTACACGACGCTCTTCCGATCT**NNNNNATGGCCACCAACAACCAGAGC** |
| 5_rev_index1 | CAAGCAGAAGACGGCATACGAGAT**ATCAGT**GTGACTGGAGTTCAGACGTGTGCTCTTCCGATCTNNNNNGAGGTTGTACGTGCCGGTCGC |
| 4_Forward adapter | AATGATACGGCGACCACCGAGATCTACACTCTTTCCCTACACGACGCTCTTCCGATCT**NNNNNACCTACCTGGCGGTGACCAGA** |
| 4_rev_index1 | CAAGCAGAAGACGGCATACGAGAT**TAGTTG**GTGACTGGAGTTCAGACGTGTGCTCTTCCGATCTNNNNNTCCACGGTCGGCAGGTTGCTG |
| LS588_rev_index2 | CAAGCAGAAGACGGCATACGAGAT**ACATCG**GTGACTGGAGTTCAGACGTGTGCTCTTCCGATCTNNNNNGACCATGCCTGGAAGAACGCC |
| LS588_rev_index3 | CAAGCAGAAGACGGCATACGAGAT**GCCTAA**GTGACTGGAGTTCAGACGTGTGCTCTTCCGATCTNNNNNGACCATGCCTGGAAGAACGCC |
| LS588_rev_index4 | CAAGCAGAAGACGGCATACGAGAT**TGGTCA**GTGACTGGAGTTCAGACGTGTGCTCTTCCGATCTNNNNNGACCATGCCTGGAAGAACGCC |
| LS588_rev_index5 | CAAGCAGAAGACGGCATACGAGAT**CACTGT**GTGACTGGAGTTCAGACGTGTGCTCTTCCGATCTNNNNNGACCATGCCTGGAAGAACGCC |
| LS588_rev_index6 | CAAGCAGAAGACGGCATACGAGATATTGGCGTGACTGGAGTTCAGACGTGTGCTCTTCCGATCTNNNNNGACCATGCCTGGAAGAACGCC |
| LS588_rev_index7 | CAAGCAGAAGACGGCATACGAGAT**GATCTG**GTGACTGGAGTTCAGACGTGTGCTCTTCCGATCTNNNNNGACCATGCCTGGAAGAACGCC |
| LS588_rev_index8 | CAAGCAGAAGACGGCATACGAGAT**TCAAGT**GTGACTGGAGTTCAGACGTGTGCTCTTCCGATCTNNNNNGACCATGCCTGGAAGAACGCC |
| LS453_rev_index2 | CAAGCAGAAGACGGCATACGAGATAAGCTAGTGACTGGAGTTCAGACGTGTGCTCTTCCGATCTNNNNNGGTCCCGAATGTCACTCGCTC |
| LS453_rev_index3 | CAAGCAGAAGACGGCATACGAGATGTAGCCGTGACTGGAGTTCAGACGTGTGCTCTTCCGATCTNNNNNGGTCCCGAATGTCACTCGCTC |
| LS453_rev_index4 | CAAGCAGAAGACGGCATACGAGAT**TACAAG**GTGACTGGAGTTCAGACGTGTGCTCTTCCGATCTNNNNNGGTCCCGAATGTCACTCGCTC |
| LS453_rev_index5 | CAAGCAGAAGACGGCATACGAGAT**TTGACT**GTGACTGGAGTTCAGACGTGTGCTCTTCCGATCTNNNNNGGTCCCGAATGTCACTCGCTC |
| LS453_rev_index6 | CAAGCAGAAGACGGCATACGAGATGGAACTGTGACTGGAGTTCAGACGTGTGCTCTTCCGATCTNNNNNGGTCCCGAATGTCACTCGCTC |
| LS453_rev_index7 | CAAGCAGAAGACGGCATACGAGAT**TGACAT**GTGACTGGAGTTCAGACGTGTGCTCTTCCGATCTNNNNNGGTCCCGAATGTCACTCGCTC |
| LS453_rev_index8 | CAAGCAGAAGACGGCATACGAGAT**GGACGG**GTGACTGGAGTTCAGACGTGTGCTCTTCCGATCTNNNNNGGTCCCGAATGTCACTCGCTC |
| Anc_rev_index2 | CAAGCAGAAGACGGCATACGAGAT**GCGGAC**GTGACTGGAGTTCAGACGTGTGCTCTTCCGATCTNNNNNTAAGGCTCCCTGGCTGTTGAC |
| Anc_rev_index3 | CAAGCAGAAGACGGCATACGAGAT**TTTCAC**GTGACTGGAGTTCAGACGTGTGCTCTTCCGATCTNNNNNTAAGGCTCCCTGGCTGTTGAC |
| Anc_rev_index4 | CAAGCAGAAGACGGCATACGAGAT**GGCCAC**GTGACTGGAGTTCAGACGTGTGCTCTTCCGATCTNNNNNTAAGGCTCCCTGGCTGTTGAC |
| Anc_rev_index5 | CAAGCAGAAGACGGCATACGAGAT**CGAAAC**GTGACTGGAGTTCAGACGTGTGCTCTTCCGATCTNNNNNTAAGGCTCCCTGGCTGTTGAC |
| Anc_rev_index6 | CAAGCAGAAGACGGCATACGAGATCGTACGGTGACTGGAGTTCAGACGTGTGCTCTTCCGATCTNNNNNTAAGGCTCCCTGGCTGTTGAC |
| Anc_rev_index7 | CAAGCAGAAGACGGCATACGAGAT**CCACTC**GTGACTGGAGTTCAGACGTGTGCTCTTCCGATCTNNNNNTAAGGCTCCCTGGCTGTTGAC |
| Anc_rev_index8 | CAAGCAGAAGACGGCATACGAGAT**GCTACC**GTGACTGGAGTTCAGACGTGTGCTCTTCCGATCTNNNNNTAAGGCTCCCTGGCTGTTGAC |
| 5_rev_index2 | CAAGCAGAAGACGGCATACGAGAT**GCTCAT**GTGACTGGAGTTCAGACGTGTGCTCTTCCGATCTNNNNNGAGGTTGTACGTGCCGGTCGC |
| 5_rev_index3 | CAAGCAGAAGACGGCATACGAGAT**AGGAAT**GTGACTGGAGTTCAGACGTGTGCTCTTCCGATCTNNNNNGAGGTTGTACGTGCCGGTCGC |
| 5_rev_index4 | CAAGCAGAAGACGGCATACGAGAT**CTTTTG**GTGACTGGAGTTCAGACGTGTGCTCTTCCGATCTNNNNNGAGGTTGTACGTGCCGGTCGC |
| 4_rev_index2 | CAAGCAGAAGACGGCATACGAGAT**CCGGTG**GTGACTGGAGTTCAGACGTGTGCTCTTCCGATCTNNNNNTCCACGGTCGGCAGGTTGCTG |
| 4_rev_index3 | CAAGCAGAAGACGGCATACGAGAT**ATCGTG**GTGACTGGAGTTCAGACGTGTGCTCTTCCGATCTNNNNNTCCACGGTCGGCAGGTTGCTG |
| 4_rev_index4 | CAAGCAGAAGACGGCATACGAGAT**TGAGTG**GTGACTGGAGTTCAGACGTGTGCTCTTCCGATCTNNNNNTCCACGGTCGGCAGGTTGCTG |
| 2-7mer Forward adapter | AATGATACGGCGACCACCGAGATCTACACTCTTTCCCTACACGACGCTCTTCCGATCT**NNNNNTCTACCAACCTCCAGAGAGG** |
| rev_index1 | CAAGCAGAAGACGGCATACGAGAT**CGTGAT**GTGACTGGAGTTCAGACGTGTGCTCTTCCGATCTNNNNNGTTGACATCTGCGGTAGCTG |
| rev_index2 | CAAGCAGAAGACGGCATACGAGAT**ACATCG**GTGACTGGAGTTCAGACGTGTGCTCTTCCGATCTNNNNNGTTGACATCTGCGGTAGCTG |
| rev_index3 | CAAGCAGAAGACGGCATACGAGAT**GCCTAA**GTGACTGGAGTTCAGACGTGTGCTCTTCCGATCTNNNNNGTTGACATCTGCGGTAGCTG |
| rev_index4 | CAAGCAGAAGACGGCATACGAGAT**TGGTCA**GTGACTGGAGTTCAGACGTGTGCTCTTCCGATCTNNNNNGTTGACATCTGCGGTAGCTG |
| rev_index5 | CAAGCAGAAGACGGCATACGAGAT**CACTGT**GTGACTGGAGTTCAGACGTGTGCTCTTCCGATCTNNNNNGTTGACATCTGCGGTAGCTG |
| rev_index6 | CAAGCAGAAGACGGCATACGAGATATTGGCGTGACTGGAGTTCAGACGTGTGCTCTTCCGATCTNNNNNGTTGACATCTGCGGTAGCTG |
| rev_index7 | CAAGCAGAAGACGGCATACGAGAT**GATCTG**GTGACTGGAGTTCAGACGTGTGCTCTTCCGATCTNNNNNGTTGACATCTGCGGTAGCTG |
| rev_index8 | CAAGCAGAAGACGGCATACGAGAT**TCAAGT**GTGACTGGAGTTCAGACGTGTGCTCTTCCGATCTNNNNNGTTGACATCTGCGGTAGCTG |
| rev_index9 | CAAGCAGAAGACGGCATACGAGAT**CTGATC**GTGACTGGAGTTCAGACGTGTGCTCTTCCGATCTNNNNNGTTGACATCTGCGGTAGCTG |
| rev_index10 | CAAGCAGAAGACGGCATACGAGAT**AAGCTA**GTGACTGGAGTTCAGACGTGTGCTCTTCCGATCTNNNNNGTTGACATCTGCGGTAGCTG |
| rev_index11 | CAAGCAGAAGACGGCATACGAGAT**GTAGCC**GTGACTGGAGTTCAGACGTGTGCTCTTCCGATCTNNNNNGTTGACATCTGCGGTAGCTG |
| rev_index12 | CAAGCAGAAGACGGCATACGAGAT**TACAAG**GTGACTGGAGTTCAGACGTGTGCTCTTCCGATCTNNNNNGTTGACATCTGCGGTAGCTG |
| F_adapter_GFPBC | AATGATACGGCGACCACCGAGATCTACACTCTTTCCCTACACGACGCTCTTCCGATCTNNNNNGGCCATCAAGCTTATCGATACC |
| R_adapter_GFPBC | CAAGCAGAAGACGGCATACGAGATCGTGATGTGACTGGAGTTCAGACGTGTGCTCTTCCGATCTNNNNNCTGATCAGCGAGCTCTAGTCG |
| NHP GAPD F | TGCACCACCAACTGCTTAGC |
| NHP GAPD R | GGCATGGACTGTGGTCATGAG |
| K9 GAPDH F | TGTCCCCACCCCCAATGTATC |
| K9 GAPDH R | CTCCGATGCCTGCTTCACTACCTT |

**Supplemental Table 2.** Primers used in the study.
